# Fine-tuning sequence to function deep learning models on large-scale proteomic data improves the accuracy of variant effect prediction

**DOI:** 10.1101/2025.09.26.678908

**Authors:** Eduarda Vaz, Lena Wang, Jake Galvin, Rebecca Keener, Alexis Battle

## Abstract

Fine-tuning sequence to function models has shown promise for variant effect prediction, but accuracy and generalization to unseen genes and unseen individuals remains a standing challenge. We fine-tuned Borzoi on 54,219 individuals and 2,923 circulating plasma proteins from the UK Biobank Plasma Proteomic Project. Across 150 single-gene models where the genes had a range of cis-heritability we observed that the fine-tuned Borzoi model improved variant effect prediction for 86% of the genes compared to an Elastic Net baseline model. We demonstrated that the improved prediction stems from increased sample size which provides tremendous amounts of rare genetic variants (MAF < 0.01) to the training data. Masking rare and uncommon variants nullified improved performance of fine-tuned Borzoi and we showed that fine-tuned Borzoi highly weights rare variants (MAF < 0.01) while the Elastic Net model highly weights common variants (MAF > 0.05) that are enriched for regulatory regions. We evaluated the generalizability of our model on a fine-tuned Borzoi model trained jointly on varying numbers of genes and observed that these models consistently outperform the pre-trained Borzoi model, the single-gene models yield more accurate results. Together this work demonstrates the importance of including larger sample sizes and rare variants in sequence to function models for variant effect prediction and demonstrates feasibility that these models are capable of highly accurate variant effect prediction.

## Introduction

Predicting the impact of genetic variation has been a key question in genomics for many years. Linear models leveraging paired genome sequencing and multiomic datasets to infer quantitative trait loci (QTLs) and their subsequent colocalization with genome-wide association study (GWAS) signals have provided insight into the impact of genetic variants on gene regulation and how that underlies complex traits^1,2^. Examining distinct data types^3^, diverse cellular contexts^2^, and increasing cohort size^4^ has led to the identification of a QTL for every human gene and together account for a greater proportion of GWAS signals. Despite these accomplishments, many GWAS signals do not colocalize with known QTLs and furthermore, these approaches are limited to common genetic variants, excluding the majority of variants that might be relevant for personalized medicine.

Sequence to function deep learning models have shown promise with variant effect prediction using strategies parallel to those used in linear models. Incorporating diverse data modalities or cellular contexts improved prediction of relevant cell types, functional regulatory elements, and the impact of variants on transcript splicing or polyadenylation, suggesting that these models have successfully learned aspects of regulatory grammar^5–7^. However, these efforts could not account for inter-individual variation as they were trained on the reference genome alone and often invert the directionality of the variant effect^8,9^.

Recently several studies fine-tuned models with the hope of retaining learned regulatory grammar patterns and gaining understanding of the impact of inter-individual genetic variation on gene regulation. Two efforts fine-tuned a pre-trained Enformer model with paired genome sequencing and RNA-sequencing data for around 1,000 individuals and used a contrastive loss function to promote accurate prediction of expression and highlight subtle differences in expression between individuals^10,11^. These models were able to accurately predict the magnitude and direction of effect of variants on gene expression for genes included in the training set but failed to generalize to unseen genes and individuals. Further, the multi-gene Performer model fine-tuned across 1,000 genes and 1,000 individuals was still unable to generalize to unseen genes and individuals^10^, suggesting that additional inter-individual data rather than inter-gene data was required. Indeed, p-SAGE-net, a lightweight model trained solely on RNA-seq data and personal genetic variation, performed well on unseen individuals, though generalizing to unseen genes and unseen individuals remained challenging^12^. These findings suggest that as with linear modeling approaches, increasing the sample size of the training data may further improve model performance.

In this study we examined the impact of sample size on model performance by fine-tuning the Borzoi mode^l6^ using plasma proteomics data for 54,219 individuals and 2,923 circulating plasma proteins from the UK Biobank Pharma Proteomics Project (UKB-PPP)^13^. We showed that increasing sample size improved model performance for some genes compared to a linear model due to increased variants considered by the model and non-linear effects of the variants on protein expression. Together these analyses demonstrate that delivering on the promise of deep learning models in precision medicine will require larger datasets and model architectures designed to leverage personalized genome sequences.

## Results

### Fine-tuned Borzoi model outperforms linear baselines for protein level prediction

The UKB-PPP dataset used Olink PEA technology to generate protein expression level estimates of 2,923 proteins in 54,219 participants. Due to the fact that the Olink assay relies on antibody binding to proteins to estimate protein levels, the estimates can be affected by cross-reactivity and epitope-binding artifacts. To reduce such biases, for every gene we excluded all samples with missense variants in the gene body. We used Elastic Net^14^ to estimate the heritability for each gene for which protein expression levels were available and identified 150 genes spanning a range of cis-heritabilities (0.02 to 0.77) and participants (5,386 to 43,190 samples). For each person each single-nucleotide variant (SNV) within the 49,152 bp region centered on the transcription start site (TSS) for each gene was extracted from Whole Genome Sequencing (WGS) data and paired with the corresponding protein expression levels. For each gene, samples were split into 60% training, 20% validation, and 20% testing sets. We developed Hottools to minimize computational time and scalability for one-hot encoding the SNVs compared to bcftools (**Methods**). Building on the Borzoi architecture, we replaced the output head with one that integrates sequence-derived embeddings with covariates (sex, age, BMI, genotype PCs, expression PCs and eGFR) through a small multi-layer perceptron to predict protein expression as a single scalar (**Fig. 1a, Supplementary Fig. 1a**). During training, both the backbone and the new head were updated jointly. This design aims to preserve the knowledge of regulatory elements while extending its capabilities to predict variant effects associated with individual protein expression. To account for differences between data types used to train the original Borzoi model (RNA-seq, CAGE, DNase-seq, ATAC-seq, and ChIP-seq) and the proteomic data used here, we added a prediction head on top of the pre-trained trunk. In the pre-trained setting, we evaluated the trunk with its original weights and our newly added head with trained weights. In the fine-tuned setting, both the trunk and the new head were jointly fine-tuned, preserving the modified architecture while allowing all parameters to adapt to the proteomics data. We quantified the L2 difference between pre-trained and fine-tuned weights across all layers to identify how this architecture impacted the weights. On average, the bias weights in the final convolution show the largest changes (**Supplementary Fig. 1**). We evaluated single-gene models on held-out individuals (1,077 to 8,238 samples) by assessing prediction accuracy with two metrics: the coefficient of determination (R^2^), which quantifies the proportion of variance explained, and the Pearson’s correlation coefficient (PCC), which measures the linear correlation between observed and predicted expression. These metrics were compared across a pre-trained Borzoi model, a fine-tuned Borzoi model, and an Elastic Net model, which functioned as a linear baseline. The fine-tuned model consistently outperformed the pre-trained model across both metrics (**Fig. 1b; Supplementary Fig. 2**), achieving a mean R^2^ of 0.62 and mean PCC of 0.78, compared to 0.22 and 0.58, respectively. Elastic Net performed slightly worse with a mean R^2^ of 0.60 and mean PCC of 0.77. Furthermore, 130 out of 150 genes (86%) with a wide range of heritability estimates had a higher PCC in the fine-tuned model compared to the Elastic Net model (**Fig. 1b,c; Supplementary Fig. 2a**). Furthermore, some genes with lower estimated heritability (HGFAC) demonstrated greater improved performance than genes with larger heritability estimates, suggesting that factors other than cis-heritability may impact model performance. *PSCA* is a highly heritable gene (cis-heritability = 0.69) with samples from 39,616 individuals in UKB-PPP. For this gene, the fine-tuned Borzoi model substantially outperformed both the pre-trained and the linear baseline (PCC = 0.97 versus 0.19 and 0.88, respectively; **Fig. 1d**). We further compared fine-tuned Enformer ^5^ to fine-tuned Borzoi, as well as fine-tuned Borzoi employing alternative head architectures without covariates. The fine-tuned Borzoi with integrated covariates demonstrated the best model performance of all models evaluated (**Supplementary Fig. 2a, b**). Notably, fine-tuned Borzoi model performance did not show a direct correlation with gene-specific sample size (**Supplementary Fig. 2b)**. The poor accuracy of the pre-trained Borzoi model likely reflects its limited exposure to personal genomic variation, whereas fine-tuning enhances the model’s ability to capture variant effects and accurately predict individual-level protein expression, as has been observed previously^10–12^. Together these results demonstrate that despite the absence of proteomic data in the pre-trained Borzoi model, after fine-tuning on personalized sequences the model is able to predict the impact of SNVs on protein expression levels for genes with a wide range of heritability in single-gene models.

**Fig 1.**
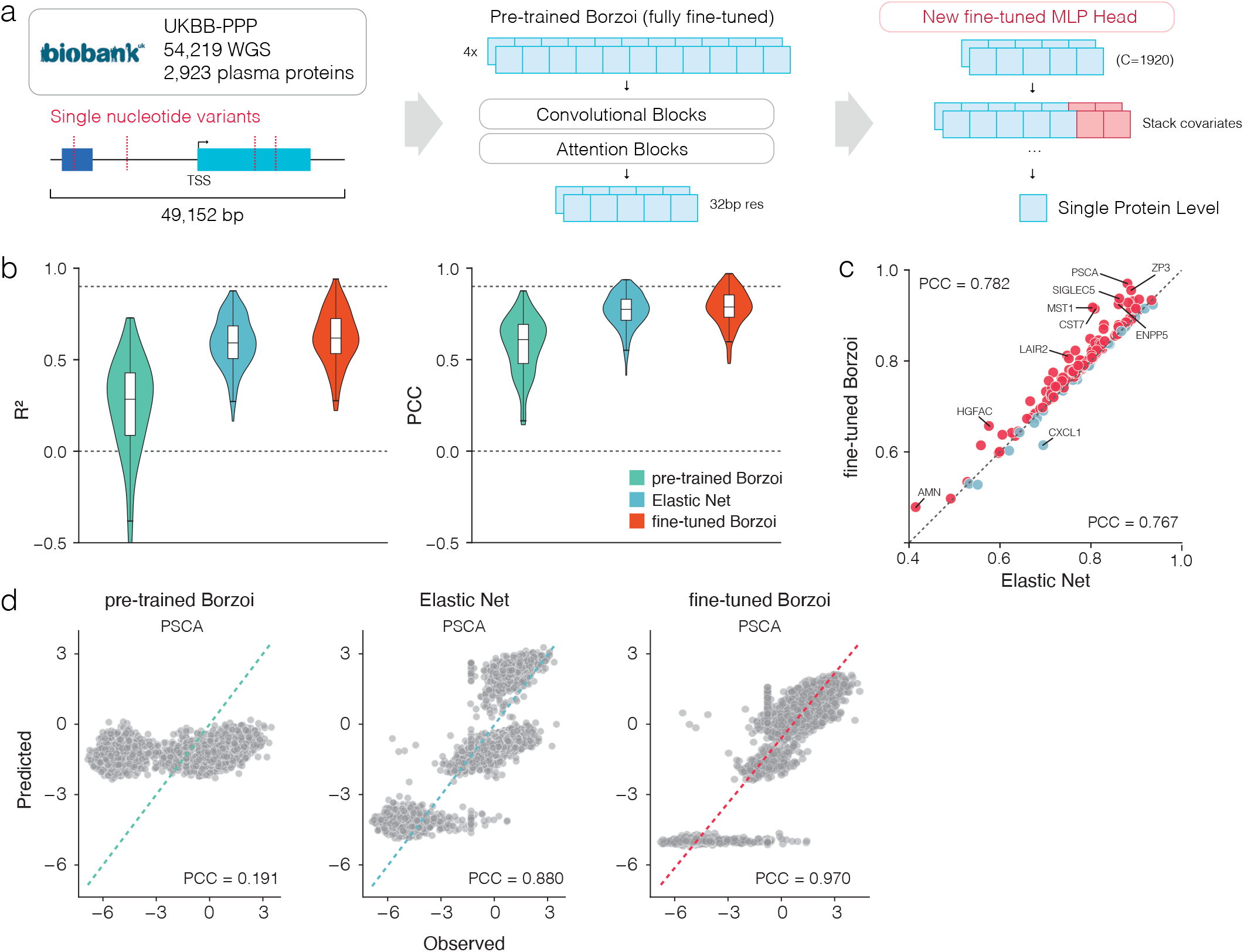
Fine-tuned Borzoi model outperforms linear baselines for protein level prediction. **a**, Overview of the UKB-PPP dataset (54,219 individuals with whole-genome sequencing and plasma proteomics for 2,923 proteins). Variants within a 49,152 bp window centered on the TSS were extracted and converted into a one-hot encoding as input to the model using Hottools. The model consists of the pre-trained Borzoi backbone with convolutional and attention blocks, followed by a new MLP head that incorporates covariates for single-protein prediction. During training, both the backbone and the new head were updated jointly. **b**, Violin plots comparing the distribution of prediction accuracy across genes on test individuals, measured by coefficient of determination (R2, left) and Pearson correlation coefficient (PCC, right), for the pre-trained Borzoi, Elastic Net, and fine-tuned Borzoi models. Each plot shows the median, interquartile range and 1.5x interquartile range as whiskers. **c**, Gene-level Pearson correlations between predicted and observed expression on test individuals, using either fine-tuned Borzoi (mean PCC=0.782) or Elastic Net (mean PCC=0.767). Each point represents a gene; red points indicate genes where fine-tuned Borzoi outperformed Elastic Net, and blue points indicate the opposite. **d**, Example predictions for the PSCA gene. Scatter plots show observed versus predicted protein expression for the pre-trained Borzoi (mean PCC=0.19), Elastic Net (mean PCC=0.88) and fine-tuned Borzoi models (mean PCC=0.97). Each point represents an individual from the test set.

### Large sample size increases rare variant representation which improves model performance

We hypothesized that three factors may contribute to improved fine-tuned Borzoi performance: cis-heritability estimates for the gene, increased sample size, and the minor allele frequency (MAF) of genetic variants. Having already demonstrated that fine-tuned Borzoi outperforms an Elastic Net model on genes with a wide variety of cis-heritability estimates (**Fig. 1c**), we next focused on the impact of sample size and the extent of genetic variation on model performance. While common variants (MAF > 0.01) are typically well represented in relatively few individuals, as the sample size per gene increased the number of rare variants (MAF < 0.01) increased substantially (**Fig. 2a**). To assess the impact of varying sample size, we trained single-gene models on downsampled cohorts of 200, 500, 1,000 and 5,000 individuals for the same set of 150 genes used in our single-gene models. For comparison, we trained Elastic Net models on the same subsets. Prediction accuracy, measured by PCC, declined with smaller sample sizes similarly between both fine-tuned Borzoi and Elastic Net models, with a loss of ∼0.1 between the full model and 200 individual cohorts in both cases (**Fig. 2b**). Across all genes the fine-tuned Borzoi model steadily improved and ultimately outperformed Elastic Net in the full sample size setting (**Fig. 2c**).

**Fig 2.**
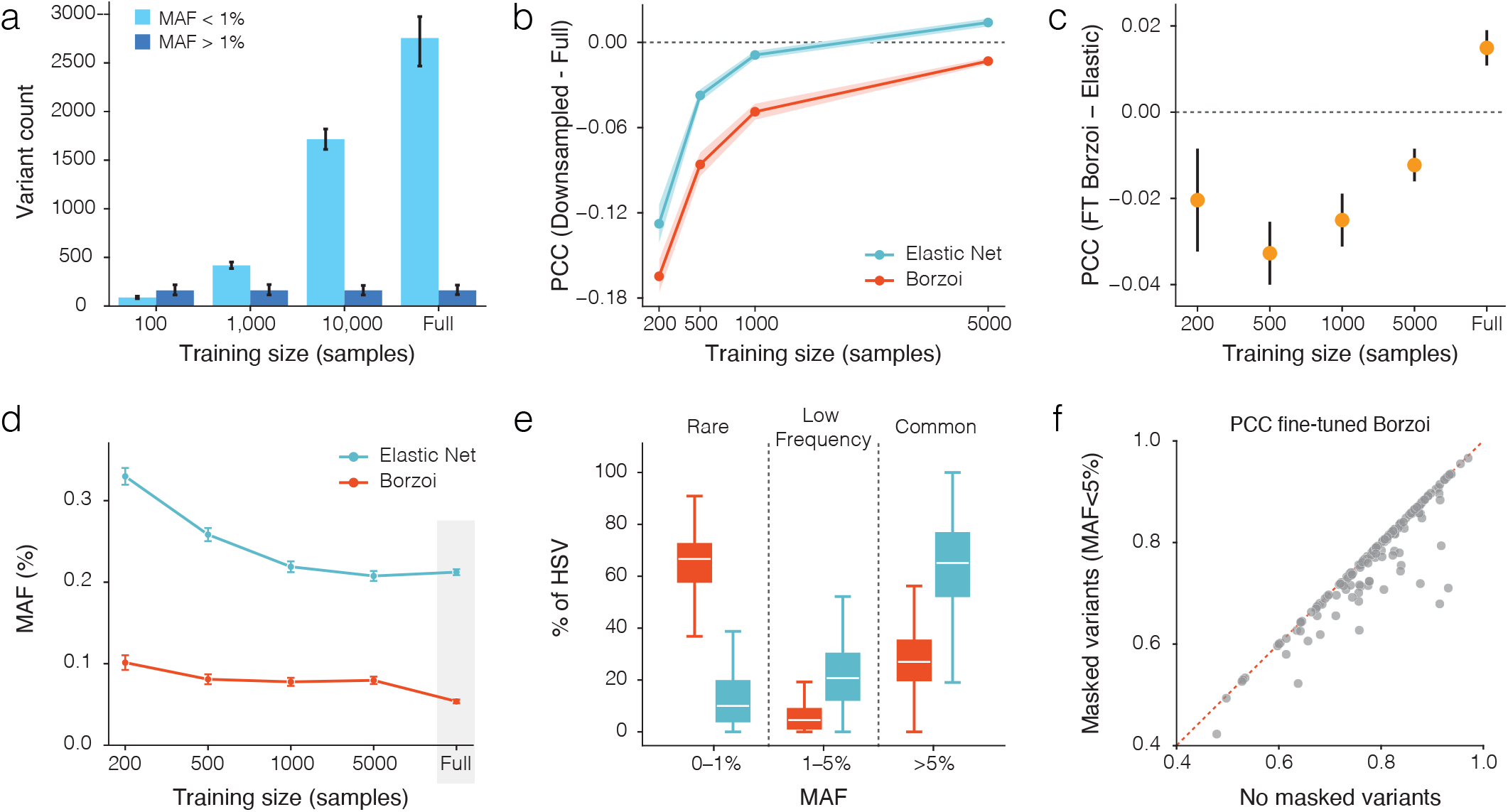
Large sample size increases rare variant representation which improves model performance. **a**, The number of variants within the input sequence (49kb region centered on the TSS) per gene at different training sample sizes, stratified by minor allele frequency (MAF>1% vs. MAF<1%). Bars show the mean and error bars the 95% confidence interval. **b**, Effect of training cohort on prediction accuracy. Single-gene fine-tuned Borzoi and Elastic Net models were trained on downsampled cohorts of 200, 500, 1,000, or 5,000 individuals for the same set of 150 genes. Prediction accuracy measured by Pearson correlation is shown relative to the full-cohort model evaluated on the same test individuals. Lines represent the mean across genes and shaded areas represent the 95% confidence interval. **c**, Difference in PCC between fine-tuned Borzoi and Elastic Net across training sizes, showing an increasing fine-tuned Borzoi advantage with larger cohorts. Points represent the mean across genes and error bars the 95% confidence interval. **d**, Mean MAF of HSVs identified by each model across training sizes. ISM was applied across the 49 kb sequence window to compute per-variant effect scores. HSVs were defined as the top n variants ranked by absolute ISM score, where n was matched to the number of non-zero Elastic Net coefficients. Fine-tuned Borzoi consistently prioritized variants with lower MAF than Elastic Net, with mean MAF decreasing further as training size increased. Points represent the mean across genes and error bars represent the 95% confidence interval. The shaded region on the full sample size is explored further in panel e. **e**, Distribution of HSVs stratified by MAF (rare: 0–1%; low-frequency: 1–5%; common: >5%) for fine-tuned Borzoi and Elastic Net in the full cohort setting. **f**, Effect of masking rare variants on prediction accuracy. Fine-tuned Borzoi models trained on the full dataset were evaluated on test individuals using either the full personal genome or a rare-masked genome in which all variants with MAF < 0.05 were substituted with the reference allele. Masking substantially reduced prediction accuracy measured by Pearson correlation, indicating that fine-tuned Borzoi relies on these variants to capture inter-individual differences.

Next, we examined which variants contributed most to model performance. We performed *in silico* mutagenesis (ISM) by systematically mutating each position in the input sequence and measuring the change in predicted expression between the mutated and reference sequence. This yielded a per-variant ISM effect score across the entire sequence. We then defined high-scoring variants (HSVs) as the top *n* variants ranked by absolute ISM score, with *n* matching the number of non-zero Elastic Net coefficients to allow direct comparison between the models. Results suggest that the fine-tuned Borzoi model prioritizes rarer variants, whereas Elastic Net prioritizes more common variants. The average MAF of HSVs was consistently lower for the fine-tuned Borzoi model than for Elastic Net, and decreased further with larger training cohorts (from 0.1 to 0.05 vs. 0.3 to 0.2). This indicates that, as sample size grows, the fine-tuned model increasingly relies on rare variants, reflecting their greater impact on capturing inter-individual expression differences in large cohorts (**Fig. 2d**). Indeed, in the full cohort setting, rare variants (MAF < 0.01) dominated the HSV set for fine-tuned Borzoi (67%), whereas in the Elastic Net common variants (MAF > 0.05) were most frequent (65%) (**Fig. 2e**). To isolate the effect of rare variants when making predictions, we evaluated fine-tuned Borzoi models trained on the full set of samples under two input settings: (1) each individual’s full personal genome and (2) a rare variant-masked genome in which all rare/low-frequency variants (MAF < 0.05) were replaced by the reference allele. Because models and training data were unchanged, any performance difference reflects the contribution of these variants. Masking variants with MAF < 0.01 had a modest effect (**Supplementary Fig. 3a**), whereas masking variants with MAF < 0.05 substantially reduced prediction accuracy (**Fig. 2f**), indicating that Borzoi relies on rare and low-frequency variants to explain inter-individual differences.

### The fine-tuned Borzoi model prioritizes functional variants that influence gene expression

To deliver on the promise of deep learning sequence to function models in clinical genomics these models must be able to prioritize functional variants. We examined whether the fine-tuned Borzoi model was capable of this by examining the intersection of regulatory element genomic annotations with ISM maps. **Fig. 3a** shows ISM maps around a rare variant (MAF < 0.001) highly scored by the fine-tuned Borzoi, which lies in an ENCODE distal enhancer-like signature region and disrupts an TEAD1 motif. In contrast, Elastic Net upweighted nearby common variant. To extend this analysis, we annotated HSVs with ChromHMM states^15^ in primary mononuclear cells from peripheral blood and observed that fine-tuned Borzoi HSVs were more enriched in active promoter and enhancer states than Elastic Net NSVs (**Fig. 3b**). Furthermore, annotations from ENCODE data tracks^16^ revealed that HSVs from the fine-tuned Borzoi model were more likely to overlap regulatory regions such as open chromatin, promoter, and enhancer regions compared to those from Elastic Net (average fraction = 0.36 vs. 0.17; **Fig. 3c**). Finally, we evaluated whether HSVs impacted transcription factor binding motifs by comparing reference versus alternate PWM log-odds using HOCOMOCO v11 PWMs^17^. Fine-tuned Borzoi models attributed higher importance to variants with a TF binding motif effect (loss/gain or |ΔS|≥0.5 weaken/strengthen) than Elastic Net (70% vs. 60%; **Fig. 3d**). To test whether models prioritized redundant signals, we compared pairwise linkage disequilibrium (LD) among HSVs within UKB WGS data. Fine-tuned Borzoi showed substantially less redundancy, with fewer HSVs in strong LD (r^2^>0.8) with each other compared to Elastic Net (19% vs. 47%), supporting the idea that the deep learning model is better at pinpointing functional variants rather than merely tagging correlated ones (**Fig. 3e**). Furthermore, the fine-tuned Borzoi model identified HSVs up to 20 kb from the TSS (the maximum input length provided), demonstrating its ability to capture distal regulatory elements^6^, while simultaneously prioritizing a higher percentage of variants near the TSS compared to Elastic Net (**Supplementary Fig. 3a**). Finally, we evaluated whether the fine-tuned models could capture the direction and magnitude of effects for SuSiE fine-mapped pQTLs^13^. For each fine-mapped variant, we substituted the alternate allele into the reference sequence and measured the predicted change in expression relative to the unmodified reference. The fine-tuned Borzoi achieved a Spearman correlation of 0.58 between predicted and observed effect sizes, indicating strong agreement with fine-mapped pQTLs (**Fig. 3f**). Together, these analyses suggest that Borzoi fine-tuning not only preserves regulatory information encoded in the pre-trained Borzoi model but also enhances it, enabling more accurate prediction of protein expression and prioritization of functional variants with stronger biological and mechanistic evidence than linear models.

**Fig 3.**
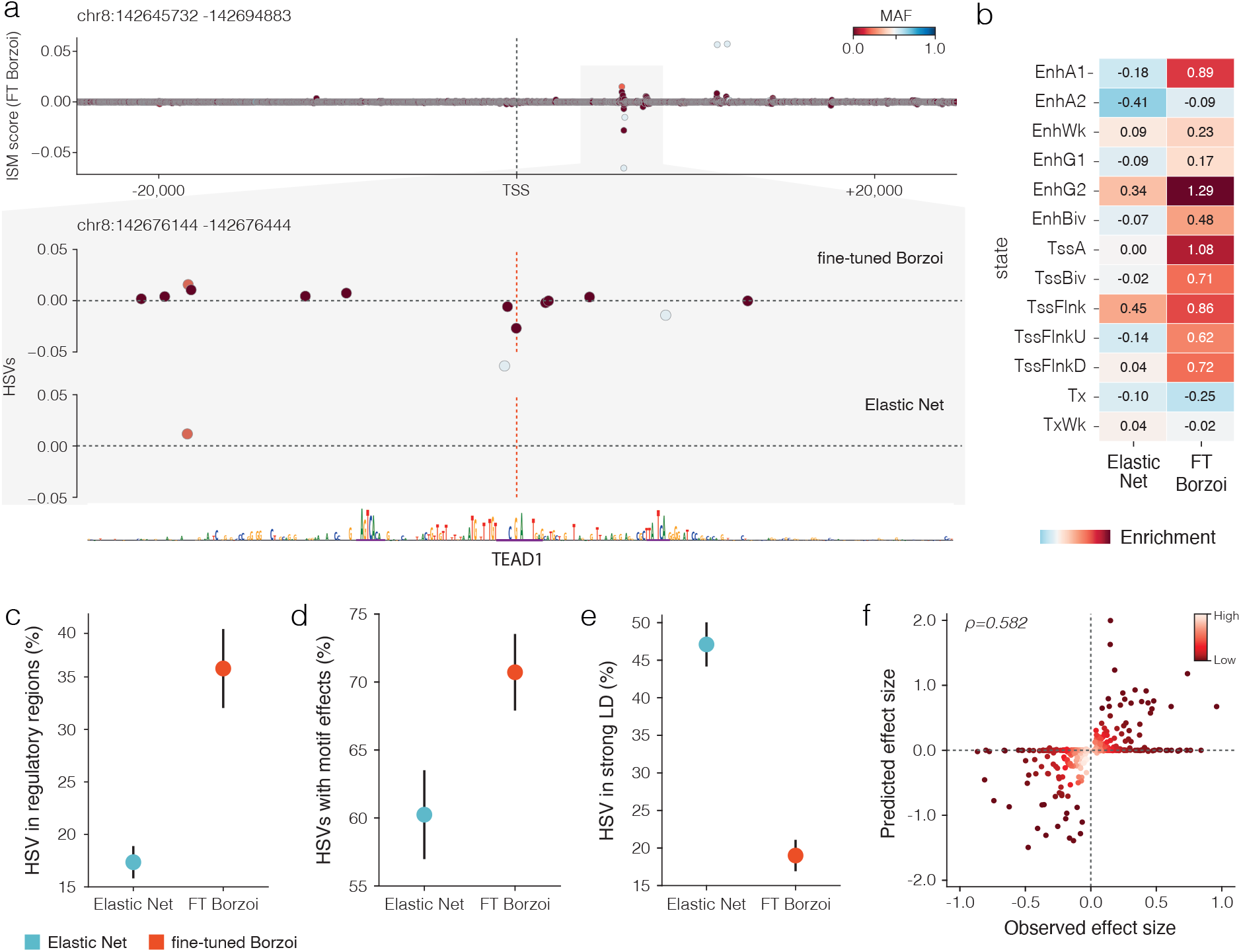
The fine-tuned Borzoi model prioritizes functional variants that influence gene expression. **a**, ISM maps around a representative rare variant (MAF<0.001) show that the fine-tuned Borzoi model assigned high importance to this variant which lies in a regulatory region (ENCODE dELS) that disrupts an TEAD1 motif, whereas Elastic Net instead upweighted nearby common variant. **b**, Enrichment of HSVs from Elastic Net and fine-tuned Borzoi (FT Borzoi) models across ChromHMM states (E062 Primary mononuclear cells from peripheral blood), showing stronger enrichment for active promoter and enhancer states in fine-tuned Borzoi. **c**, Fraction of HSVs overlapping regulatory regions from ENCODE annotations (open chromatin, promoter, enhancer), with fine-tuned Borzoi attributing more HSVs to regulatory elements than Elastic Net. Points represent the mean across genes and error bars represent the 95% confidence interval. **d**, Fraction of unique HSVs with any motif effects (loss/gain or |ΔS|≥0.5 weaken/strengthen).Fine-tuned Borzoi-prioritized HSVs showed a higher fraction with motif effects than Elastic Net. Points represent the mean across genes and error bars represent the 95% confidence interval. **e**, Fraction of HSVs in strong linkage disequilibrium (LD) (r2>0.8) with other HSVs, demonstrating that fine-tuned Borzoi produces fewer redundant signals than Elastic Net. Points represent the mean across genes and error bars represent the 95% confidence interval. **f**, Predicted versus observed effect sizes for SuSiE fine-mapped pQTLs (PIP > 0.9). Fine-tuned Borzoi achieved a Spearman correlation of 0.58, indicating strong agreement with fine-mapped effects and outperforming the pre-trained Borzoi. The color gradient represents the density of points in the scatter plot.

### Fine-tuning improves generalization to unseen genes compared to pre-trained model

A more challenging setting is predicting expression both across unseen genes and unseen individuals. Here, the model cannot rely on having observed the same gene or nearby regulatory motifs during training but must generalize from broader causal regulatory principles. We trained models jointly on multiple genes so that learned representations could transfer to novel genes and variants. Unlike the single-gene setup, we did not include covariates, since a shared head would impose identical covariate effects across genes, while defining gene-specific heads would prevent inference on unseen genes. In addition, the loss function excluded the mean squared error term and only used the pairwise difference component to focus training on variation across individuals rather than gene-specific expression levels. We evaluated multi-gene fine-tuned models across different training set sizes, varying the number of individuals (2,000-10,000) and genes (100, 200, and 300). For the test set of unseen genes, the corresponding genomic regions were also not used during the training of the pre-trained Borzoi model, ensuring that evaluation was performed entirely on novel loci. On seen genes and unseen individuals, fine-tuned models trained on multiple genes did not improve performance over single-gene models, suggesting that the model does not gain additional gene-specific information simply from exposure to other genes. Although larger sample sizes substantially improved prediction accuracy (**Fig. 4a, b**). On unseen individuals and unseen genes the multi-gene models consistently explained more inter-individual variability than the pre-trained Borzoi model trained only on the reference genome (**Fig. 4c, d**). This contrasts with previous work, where fine-tuned models failed to generalize to alleles in unseen genes and performed worse than the pre-trained Borzoi^10–12^. In our setting, performance generally improved as more individuals were included in training (**Fig. 4c, d**). Notably, we observed that increasing the number of training genes or individuals does not always yield better generalization. This may reflect overfitting to gene-specific features, bias toward dominant patterns, or catastrophic forgetting, where useful representations are lost due to conflicting training signals. While this work provides evidence that sequence to function deep learning models may feasibly generalize to unseen genes and unseen individuals, accurate prediction in this context remains a key challenge, likely requiring stronger regularization or architectural modifications.

**Fig 4.**
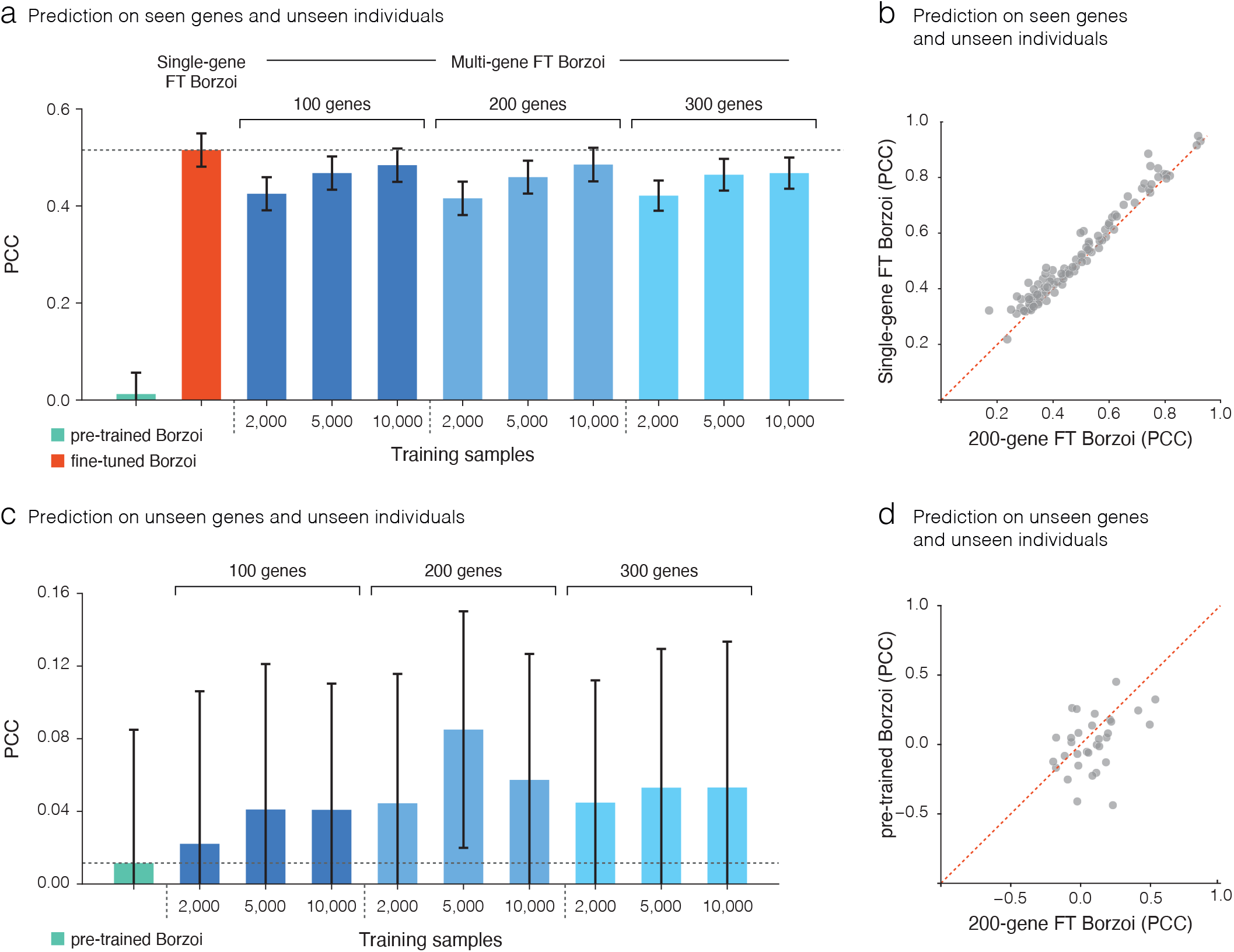
Fine-tuning improves generalization to unseen genes compared to pre-trained model. **a**, Prediction accuracy on seen genes and unseen individuals measured by Pearson correlation. Fine--tuned single-gene models outperformed the pre-trained Borzoi. Training on multiple genes (100, 200, or 300) with 2,000–10,000 individuals (blue shades) achieved comparable accuracy to single-gene models, suggesting that the model does not gain additional gene-specific information simply from exposure to other genes. Bars show the mean across test genes and error bars the 95% confidence interval. **b**, Gene-level comparison of prediction accuracy between single-gene and multi-gene (200 genes and 10,000 individuals) models on seen genes and unseen individuals. **c**, Prediction accuracy on unseen genes and unseen individuals. Multi-gene fine-tuned Borzoi models consistently explained more inter--individual variability than the pre-trained Borzoi trained only on the reference genome, with performance improving as more individuals were included in training. Notably, performance peaked at an intermediate setting (200 genes, 5,000 individuals), suggesting an optional point between generalization to unseen genes and the number of samples and genes used during training. Bars show the mean across test genes and error bars represent the 95% confidence interval. **d**, Gene-level comparison of prediction accuracy between single-gene and multi-gene (200 genes and 5,000 individuals) models on unseen genes.

## Discussion

Deep learning models require tremendous amounts of input data to identify complex patterns. The UKB-PPP dataset consists of 54,219 individuals which is approximately 50x those previously used in similar studies and we demonstrated that increased sample size directly contributed to improved model performance on unseen genes and individuals. Previous studies fine-tuning Enformer have shown the importance of common genetic variants for predicting genes with high heritability and describe a plateau effect on model performance when increasing the number of genes used in a multi-gene model^10,11^. Borzoi was previously reported to have higher accuracy in predicting the effects of fine-mapped eQTLs^6^ and our increased model performance may be in part due to our choice to fine-tune the Borzoi model instead of the Enformer model. We demonstrated the fine-tuned Borzoi model upweights rare and low frequency variants (MAF < 0.05) that are enriched for functional annotations. Increasing the sample size and thus the number of rare variants overall improved single-gene model performance compared to an Elastic Net linear model. A multi-gene fine-tuned Borzoi model trained on varying numbers of genes also suggested that model performance increased with more training samples when evaluated on unseen individuals. However, none of the multi-gene models demonstrated high accuracy on unseen genes and unseen individuals, leaving the generalization of fine-tuned sequence to function deep learning models a standing challenge.

While the UKB-PPP dataset has tremendous power it also presents several limitations. First, the Borzoi model training data did not include proteomic data, resulting in a slight mismatch between the training and application datasets. It is likely that including proteomic data and re-training the Borzoi model, or other models, would further improve performance after fine-tuning. Second, because of the technical nature of Olink proteomic estimates we removed individuals with any missense variant within the gene body to ensure the protein expression estimates were not confounded by epitope binding artifacts. This reduced our sample sizes for many genes which could likely largely be recovered if we better understood the epitopes used for Olink. Finally, this proteomic dataset was collected from plasma which hinders our ability to make context-specific insights from this data as plasma consists of proteins secreted into the blood across the human body^18^, however, this also means that this dataset comprises signatures of diverse processes. Therefore, careful examination of variant effects, for example those predicted to impact transcription factor binding sites, using this data may still produce valuable insights.

Given the promise of sequence to function deep learning models for predicting variant effects the scientific community will soon seek ways to combine greater amounts of increasingly diverse data types and in larger quantities. It is likely that some data types will be more informative for the prediction of variant effects on gene regulation and protein levels than others as more specific models have shown greater performance on this task than models trained on a wide variety of data types^19^. The resources required to generate these data at scale will be tremendous and may not be equitably distributed across institutions. Furthermore, the capacity to train increasingly large models will require technical innovation and availability of compute resources. In this context, parameter-efficient fine-tuning (PEFT) approaches offer a practical solution to adapt large models efficiently with competitive performance while updating only a small fraction of parameters^20^. Such approaches make large-scale modeling more accessible under resource constraints and will be especially valuable as genomic models continue to grow in size. A critical challenge for the field will be to innovate model architecture to continue to improve model performance at increased efficiencies.

This work contributes to the conclusion that powerful variant effect prediction deep learning models benefit from training data that model diverse molecular processes, capture diverse cellular contexts, and are produced at scale. With further innovation in model architecture, software and hardware, and the integration of data types, sequence to function deep learning models may deliver on the promise of genetics in precision medicine.

## Acknowledgements

We would like to thank Seraj Grimes, Fuchen Li, Gus Fridell and other members of the Battle lab for helpful conversations about this project. This research has been conducted using the UK Biobank Resource under Application Number 98985. A.B. was supported by NIGMS R35GM139580.

## Author contributions

E.V. participated in project conceptualization, formal analysis, investigation, methodology, data visualization, and writing. L.W. participated in formal analysis and data curation. J.G. participated in formal analysis. R.K. participated in project administration and writing. A.B. participated in project conceptualization, supervision, funding acquisition, and writing.

## Methods

### Protein expression dataset

Models were trained and evaluated using protein expression data from circulating plasma, provided by the UK Biobank Pharma Proteomics Project (UKB-PPP)^13^. The dataset comprises normalized protein expression levels (NPX) for 54,219 individuals, from which 46,595 belong to the random baseline cohort, with matched unphased whole-genome sequencing (WGS) data and measurements across 2,923 unique proteins. Details of the Olink proximity extension assay (PEA) used for proteomic profiling are described elsewhere^13^. No additional transformations were applied to the NPX values.

### Selection of proteins

We selected proteins for model training using two criteria: (1) <2,000 missing protein measurements across individuals, and (2) at least one cis-pQTL whose associated variant lay within the 49,152bp input window and was not protein-altering (frameshift, missense, start-lost, or stop-gained). Applying these criteria resulted in 688 proteins for downstream analyses.

### Variant calling and selection of individuals

A potential risk of Olink PEA is epitope binding effects, where genetic variants or post-translation modifications alter the antibody binding site, leading to inaccurate protein quantification^21,22^. To mitigate these biases, samples containing variants likely to affect protein conformation were excluded on a per-gene basis. Single-nucleotide variants (SNVs) within each protein-coding gene body were annotated with the Ensembl Variant Effect Predictor (VEP)^23^, and any individual harboring a missense, stop-gained, stop-lost, start-lost, or frameshift variant was removed for that gene. This yielded gene-specific cohorts ranging from 5,386 to 43,190 individuals. For each gene, the cohort was split into 60% training, 20% validation, and 20% testing sets across both linear and deep-learning models to ensure a fair comparison.

### Hottools: Fast one-hot encoding of individual genomic sequences from single-nucleotide variants

We present Hottools, a high-performance Python-based tool for fast reconstruction and one-hot encoding of individual genomic sequences from VCF data. Hottools extracts biallelic single-nucleotide variants (SNVs) from phased or unphased genomes and integrates them with a reference genome (GRCh38) to generate individualized sequences. It encodes both heterozygous and homozygous SNVs in a consistent one-hot format, enabling direct use in downstream machine learning pipelines. The resulting encoding represents whether an individual has no dosage (0/0; 0), half dosage (0/1; 0.5) or full dosage (1/1; 1) of any particular SNV. By default, it assigns A=[1,0,0,0], C=[0,1,0,0], G=[0,0,1,0] and T=[0,0,0,1]. Unlike existing tools such as bcftools consensus, which process samples sequentially, Hottools is optimized to process thousands of individuals simultaneously, significantly reducing runtime and improving scalability. This approach enables scalable generation of personalized sequence matrices for population-scale genomic analyses.

### Model inputs

For each individual in the UKB-PPP, personalized genomic sequences were generated using Hottools. Hottools integrates biallelic SNVs from the individual’s VCF file with a reference genome and encodes the resulting sequences in a one-hot format, capturing both homozygous and heterozygous variants. To accommodate GPU memory constraints and enable larger batch sizes during training and evaluation, we used input sequences of 49,152 base pairs (bp), shorter than Borzoi’s maximum receptive field of 524 kb. For each gene, we extracted a 49-kb one-hot encoded sequence centered on the transcription start site (TSS), as annotated in GENCODE v47.

### Fine-tuning Enformer

For fine-tuning Enformer^5^, the github.com/lucidrains/enformer-pytorch provides an easy implementation with HeadAdapterWrapper class. This wrapper was configured with num_tracks=1, post_transformer_embed=False and an identity function to produce a single scaler expression output. This bypasses the embeddings from the original human and mouse Enformer head through a new fully connected feedforward neural network to predict the new single scalar target. The model was fine-tuned end-to-end without freezing any weights. The output was a vector of length 384, corresponding to genomic bins. Only the bin that overlaps the TSS (bin 192) was used as the predicted value. This scaler was used to compute the loss function during training and compared against observed expression during evaluation.

### Fine-tuning Borzoi

The PyTorch version of Borzo^i6^ and respective pre-trained weights were obtained from github.com/Genentech/gReLU^24^. We set crop=0 and removed the task-specific heads after the last GELU, exposing a sequence embedding of shape [B, C, L]. We applied a global average pooling across the positional axis (L), concatenated a c-dimensional covariate vector (sex, age, BMI, genomic PCs, expression PCs, eGFR), and normalized the concatenated features with LayerNorm. The new head is a two-layer MLP, Linear (512), ReLu, Linear (1), yielding a single scalar prediction. All parameters (backbone and head) were fine-tuned jointly. The scalar output was used to compute the training loss against measured protein expression and for evaluation.

### Training scheme

Optimization was performed using the AdamW optimizer and a learning rate schedule that included a warm-up phase over the initial 10% of training steps, gradually increasing to 1e-4, followed by cosine decay to 0. During training the model received 49,152-bp sequences as input, with a batch size of 4 on a 24GB NIVIDIA A10 GPU with gradient accumulation of 64 steps to achieve an effective batch size of 256.

### Loss function

The loss function is composed of two terms: (1) the mean squared error between predicted and observed protein expression, and (2) the mean squared error of pairwise differences between all unique pairs of individuals within the batch, comparing the observed differences with the predicted differences. The second component serves as an inter-individual contrastive term, encouraging the model to preserve the relative differences in expression between individuals. The total loss is defined as a weighted sum of the two components, with the weighting parameter alpha set to 0.5. A closely related loss has been used in prior work^10–12^.

### Elastic Net Model and selection of proteins

Elastic net models were trained independently for each of the 688 selected proteins using a genotype matrix, where individual-level dosages (0, 1, 2) represented SNVs located within a 49,152-bp window centered on the TSS of each gene (same as for the deep learning models). We set a fixed L1 ratio of 0.5. Genes were then ranked based on the resulting coefficient of determination (R^2^), and the top 150 genes were selected for downstream analyses.

### *In silico* mutagenesis and high-scoring variants

*In silico* mutagenesis was used to measure each variant’s contribution to the fine-tuned model. For each gene, we systematically mutated every position in the 49,152 bp input sequence to each of the three non reference bases, one at a time, and measured the change in predicted expression relative to the reference sequence. Each mutated sequence was passed through the model to obtain a prediction. The three alternate predictions were averaged to compute the ISM score at that position. This yielded a per-variant effect map across the window.

To compare variant prioritization across models, we defined high-scoring variants (HSVs) per gene using matched set sizes. Specifically, we took the number of non zero coefficients from the corresponding Elastic Net model as *n*. For Elastic Net, HSVs were the variants with nonzero coefficients. For the deep models, HSVs were the top *n* positions ranked by absolute ISM effect.

### Downsampled model analysis

To isolate the effect of cohort size, we trained single-gene models on nested downsampled cohorts of 200, 500, 1,000, and 5,000 individuals for the same set of 150 genes used elsewhere. For each gene and each cohort size, we trained both a fine-tuned Borzoi model and an Elastic Net model using the same individuals within that subset. A fixed held-out test set, same individuals and size across all configurations and model classes was used for evaluation. To keep optimization comparable across cohort sizes, we reduced the effective batch size for smaller subsets so that the number of optimizer updates per training run matched that of the full-cohort model, while keeping other training hyperparameters unchanged. Downsamples were nested, individuals were added cumulatively without replacement, so differences across cohort sizes reflect the marginal effect of additional samples rather than changes in sample composition.The deep-learning models used the same 49,152-bp input window centered at each gene’s TSS as in the full model, Elastic Net used the matching genotype design matrix (0/1/2 dosages) over the same window.

### LD analysis

We quantified pairwise linkage disequilibrium (LD) among HSVs using UKB WGS data. For each gene, we computed PLINK pairwise LD (r^2^) within the 49 kb window and defined strong LD as r^2^ above 0.8. Redundancy was summarized as the fraction of HSVs with at least one HSV partner in strong LD (aggregated across genes).

### ChromHMM enrichment analysis

For each gene, the 49,152-bp input sequence was annotated using the NIH Roadmap Epigenomics Consortium ChromHMM 18-state annotations for Primary mononuclear cells from peripheral blood (E062)^15^. The chromatin states were each mapped to either an enhancer, promoter, or other regions. Observed counts were the number of HSV positions overlapping each state. Expected counts per state were based on the proportion of the sequence labeled with a given state multiplied by the total number of HSVs to get For each gene, we looked at what share of its 49,152-bp window was labeled with a given ChromHMM state. We then took that share and multiplied it by the total number of HSVs for that gene to get how many HSVs we would expect to fall in that state if HSVs were placed uniformly at random within the window. These values were then aggregated across all genes. The enrichment was calculated as the ratio of observed over expected HSVs for each state, with a high enrichment value indicating that the model was assigning more variants to a state than expected.

### Regulatory Analysis

To annotate each gene’s 49kb sequence, we used the ENCODE cCRE (Candidate Cis-Regulatory Elements) data^16^. Elements were grouped into five classes: promoter-like signatures (PLS), proximal enhancer-like signatures (pELS), distal enhancer-like signatures (dELS), DNase–H3K4me3 regions, and CTCF-only sites. Each basepair in the window was assigned to one of these five labels or none if no cCRE overlapped. For the fine-tuned Borzoi and Elastic Net models, we calculated the fraction of HSVs falling within these regulatory elements.

### Motif-impact Analysis

We evaluated whether HSVs alter TF-binding by scanning ±30 bp windows centered on each SNP with HOCOMOCO v11^17^ PWMs using FIMO (MEME Suite v5.5.2). Reference and alternate sequences were scanned with the same order-0 background (A/C/G/T frequencies from GRCh38 via fasta-get-markov) and the same prefilter (--thresh 1e-4). For each motif, we retained the top-scoring hit overlapping the SNP and computed the difference in PWM log-odds between the alternate and the reference (ΔS). Hits were called significant at q≤0.05, and we classified effects as loss (REF significant, ALT not significant), gain (ALT significant, REF not significant), weaken/strengthen (both significant with |ΔS|≥0.5), or no motif otherwise. We then collapsed to one call per variant (max |ΔS|) and reported the fraction of unique HSVs with any motif effect (loss/gain or |ΔS|≥0.5 weaken/strengthen).

### Multi-gene model and predicting unseen genes

We fine-tuned a single Borzoi model jointly across multiple genes using a shared convolutional head and no covariates. Covariates were omitted because a shared head would enforce identical covariate effects across genes, whereas per-gene heads would preclude inference on unseen genes. Each batch comprised multiple individuals from a single gene, and the loss function used only the previously defined pairwise-difference term to emphasize inter-individual variation. Optimization matched the single gene setup (AdamW with 10% warm-up to 1e-4 and cosine decay), with gradient accumulation increasing to 128 for an effective batch size of 512. We trained multiple configurations varying the number of individuals (2,000, 5,000, 10,000) and genes (100, 200, 300). For all configurations, the held-out test set was fixed, the same test individuals were used per gene, and for the unseen-gene evaluation, test genes were drawn from Borzoi’s original validation/test partitions to ensure assessment exclusively on loci not used during training.

